# Coordinated regulation of Barrier-to-Autointegration Factor (BAF) phosphorylation by protein phosphatases PP2A and PP4

**DOI:** 10.64898/2026.06.16.732553

**Authors:** Mònica Torras-Llort, Paula Escudero-Ferruz, Edit Abraham, Zoltan Lipinszki, Chong Zhang, Fernando Azorin

## Abstract

Phosphorylation is crucial for the regulation of Barrier-to-Autointegration Factor (BAF), a conserved nuclear envelope (NE)-associated protein that, among its diverse functions, binds chromatin and mediates its anchoring to the NE. In interphase, cycles of BAF phosphorylation, which inhibits chromatin binding, and dephosphorylation, which restores it, are essential for the dynamic anchoring of chromatin to the NE. During mitosis, BAF becomes highly phosphorylated and dissociates from chromatin, except for a small centromeric pool (cenBAF). Upon mitotic exit, BAF is dephosphorylated and re-associates with chromosomes. Two Ser/Thr protein phosphatases, PP2A and PP4, are known to dephosphorylate BAF, but their specific roles in regulating BAF phosphorylation are not yet fully understood. Here, we show that PP2A is the main phosphatase driving BAF dephosphorylation in interphase, whereas PP4 plays a minor role. During mitosis, both PP2A and PP4 localize to centromeres; however, only PP4 directly interacts with cenBAF, suggesting that it maintains this fraction in an unphosphorylated state. Notably, PP4’s centromeric localization depends on PP2A. At mitotic exit, PP2A dephosphorylates the large pool of mitotically phosphorylated BAF, while PP4 controls the timing of its re-association with chromatin. Together, these findings reveal a coordinated interplay between PP2A and PP4 in controlling BAF dephosphorylation and function.

## INTRODUCTION

Barrier-to-Autointegration Factor (BAF) is a highly conserved nuclear envelope (NE)-associated protein in metazoans (reviewed in (1–3)). BAF participates in diverse cellular processes, including the regulation of retroviral integration, chromatin organization, and transcription, as well as NE reassembly (NER) after mitosis and the repair of NE ruptures (reviewed in (1–5)). BAF depletion is embryonically lethal in *Caenorhabditis elegans* and *Drosophila* (6,7). In humans, BAF mutations are linked to Nestor-Guillermo progeria syndrome (NGPS), an early-onset ageing disorder (8,9), and to a dominant motor neuropathy (10).

Central to BAF function is its ability to bridge chromatin to the NE. BAF is an obligate homodimer that interacts with nuclear lamina (NL)-associated proteins, including Lamins and LEM domain proteins (11–19), and binds nucleosomal DNA in a sequence-independent manner (6,20–22). This dual interaction with DNA/nucleosomes and NL-associated proteins underlies its role in regulating chromatin-NE interactions (reviewed in (23–26)), particularly in anchoring heterochromatin to the NE (27).

Phosphorylation plays a crucial role in regulating BAF localization and function. *In vitro* studies have shown that phosphorylation inhibits BAF binding to DNA (28–32), while *in vivo*, BAF is extensively phosphorylated during mitosis coinciding with its released from chromatin (29,31,33,34). Consequently, impairing BAF phosphorylation causes its retention on mitotic chromosomes (27,29,35), indicating that BAF phosphorylation is required for chromatin dissociation. Notably, although during mitosis the majority of BAF is phosphorylated and free in solution, a small fraction remains associated with centromeric chromatin, referred to as cenBAF (36). During mitotic exit, BAF is dephosphorylated and re-associates with chromosomes in a process that drives post-mitotic NE reassembly (14,29,34,37–40). Dynamic BAF phosphorylation is also important for efficient anchoring of heterochromatin to the NE during interphase (27).

BAF phosphorylation is mediated by the vaccinia-related kinase VRK1/2 (NHK1 in *Drosophila*), which targets conserved Ser/Thr residues in its N-terminus (28,31,41). In contrast, two distinct heterodimeric protein phosphatase complexes, PP2A and PP4, have been shown to reverse this modification (34,38,42). PP2A complexes consist of an evolutionarily conserved PP2Ac catalytic subunit (Microtubule Star (MTS) in *Drosophila*), a structural A subunit, and variable B regulatory subunits involved in substrate recognition (Twins (TWS) and Widerborst (WDB) in *Drosophila*) (43–45). The PP4 holoenzyme, on the other hand, consists of a conserved PP4c catalytic subunit, a scaffolding R2 subunit, and an R3 regulatory subunit (Falafel (Flfl) in *Drosophila*) that mediates substrate recognition (46–49). How the activities of PP2A and PP4 complexes differentially regulate BAF phosphorylation and function remains unclear. In this study, we address this question. Our results indicate that PP2A acts as the major BAF phosphatase during mitosis exit and interphase, whereas PP4 function appears to be spatially restricted to centromeres, where it is recruited by the constitutive centromeric protein CENP-C (50) and directly interacts with cenBAF.

## MATERIALS AND METHODS

### DNA constructs

Plasmids YFP::BAF and YFP::BAF^T44/S55A^, expressing the indicated YFP-tagged BAF constructs under the control of the endogenous *BAF* promoter, and act5-mCherry::HP1a are described in (27). pMT[Blast]_eGFP-CENP-C-RES (resistant to RNAi) is described in (36). The N-terminally 3xFlag-tagged constructs (pMT[Hygro]_3xFlag-Flfl, pMT[Hygro]_3xFlag-MTS, pMT[Hygro]_3xFlag-WDB and pMT[Hygro]_3xFlag-TWS-RA) were placed under the control of the copper-inducible metallothionein promoter (pMT), are hygromycin-selectable, and were generated using standard PCR and Gateway cloning methods. Briefly, CDS of PP2Ac/*mts* (*microtubule star*, cDNA clone: LD26077), PP2A-B’/*wdb* (*widerborst*, cDNA clone: LD34343), PP2A-B/*tws-RA* (*twins-RA*, cDNA clone: LD12394) or PP4R3/*flfl* (*falafel*, cDNA clone: LD13350) were PCR-amplified using gateway primers (**Supplementary Table S1**) and cloned into the pDONR221 plasmid (Thermo Fisher Scientific, cat#12536017) by BP recombination. Entry clones were sequence-verified and CDS were recombined into the pMT[Hygro]_3xFlag-DEST plasmid using LR clonase. The pMT[Hygro]_3xFlag-DEST plasmid was generated in-house as follows: the ‘3xFlag-Gateway’ cassette was subloned from pAFW (DGRC, cat#1111) to the KpnI-PmeI site of the pMT/V5-His A plasmid (Thermo Fisher Scientific, cat# K4120-01) to create the pMT_3xFlag-DEST vector. The ‘HindIII-copia promoter-hygromycin resistance gene-SV40 polyA signal-Ecl136II’ cassette was subloned from the pCoHygro plasmid (Thermo Fisher Sceintific, cat# K4120-01) into the KspAI site of pMT_3xFlag-DEST to generate the pMT[Hygro]_3xFlag-DEST plasmid.

### Antibodies

Rabbit polyclonal αBAF and rat and rabbit polyclonal αCENP-C antibodies are described in (27,36). Rat polyclonal αFlfl and rabbit polyclonal αPP4c are described in (50), rabbit αTWS was a gift from Dr. Vincent Archambault, and guinea pig αWDB was a gift from Dr. Amita Sehgal. The rest of antibodies used were commercially available: mouse monoclonal αMTS (BD-Transduction Laboratories TM, 610555), mouse monoclonal αLaminDm0 (DSHB ADL67.10), rabbit polyclonal αGFP (Invitrogen A11122), mouse polyclonal αGFP (Roche 118144600), mouse monoclonal αFlag M2 (Merck Millipore F3165), rabbit polyclonal αFlag (Merck life science F7425), rabbit polyclonal αH3 (Cell Signaling 9715), and mouse monoclonal αbetaTubulin (MAB3408 Merck Millipore).

### Stable *Drosophila* S2 cell lines

Schneider’s *Drosophila* Line 2 [D. Mel. (2), SL2] (ATCC® CRL-1963™) were maintained in complete serum-free medium (cSFM) consisting of Insectagro® Medium (Corning, 13-402-CV) supplemented with 2 mM L-glutamine (Biosera XC-T1755/100) and 1x PenStrep (Biosera XC-A4122/100). Double-stable cells lines co-expressing GFP::CENP-C-RES and N-terminally 3xFlag-tagged Flfl, TWS-RA, WDB or MTS under control of the copper-inducible pMT promoter were generated by standard procedures. Briefly, D. Mel. (2) cells were co-transfected in 6-well plates with 3 µg of pMT[Blast]_eGFP-CENP-C-RES and 3 µg of each pMT[Hygro]_3×Flag construct using Cellfectin II transfection reagent in cSFM, according to the manufacturer’s instructions (Thermo Fisher Scientific, 10362100). Three hours post-transfection, the medium was replaced with fresh cSFM, and cells were incubated for 3 days at 25 °C. Selection was then initiated by replacing the medium with cSFM containing 25 µg/mL blasticidin S (Corning, cat#30-100-RB) and 300 µg/mL hygromycin B (Serva, cat#25965), and cells were cultured for an additional 3 days at 25 °C. Dual antibiotic selection was maintained according to standard procedures until passage 5, at which point cells were cryopreserved and used for experiments. Co-expression of eGFP::CENP-C-RES and 3×Flag::Flfl, 3×Flag::Mts, 3×Flag::Wdb, or 3×Flag::Tws-RA was confirmed by western blotting using αGFP, αFlag M2 and αCENP-C antibodies in cells induced with 1 mM CuSO_4_ for 24 hours **(Supplementary Figure S1).**

### RNAi knockdown experiments

RNAi-mediated knockdown experiments were performed as previously described (36). Briefly, amplicons were generated using the primers listed in **Supplementary Table S1** and synthesized using the MEGAscript T7 Transcription Kit (Ambion AM1334). For knockdowns, S2 cells were grown to a density of 5×10^5^ cells/mL and incubated with 30-40 μg of dsRNA. After 3 days, cells were diluted 1:2 and treated with a second 30 μg dose of dsRNA for an additional 4 days. For WDB knockdown, a combination of two distinct amplicons was employed, consisting of 20 μg each of dsWDB1 and dsWDB2 per dose. For MTS knockdown a single 35 μg dose for 3 days was used. The efficiency of protein depletion was determined by WB (**Supplementary Figure S2**).

### Western blot analysis

Western blot analyses were performed according to standard procedures using the following antibody dilutions: αBAF (1:2500), αFlfl (1:10000), αMTS (1:1000), αWDB (1:2000), αPP4c (1:5000), αH3 (1:2500), αTubulin (1:5000), αCENP-C (1:3000), mouse αGFP (1:2000), mouse αFlag (1:2500) and αLaminDm0 (1:5000).

### Okadaic acid treatment and analysis of BAF phosphorylation

To analyze the phosphorylation status of BAF, control and depleted cells were incubated for 3 hours with either DMSO (vehicle control) or 20 nM Okadaic acid (OA). Following treatment, cells were washed and allowed to recover for 3, 6, 21, or 30 hours prior to harvest. Total protein was extracted using a lysis buffer consisting of 50 mM Tris-HCl (pH 8.0), 150 mM NaCl, 10% glycerol, 1% NP-40, and 0.1% SDS. To maintain protein integrity and phosphorylation states, the buffer was supplemented with 1 mM PMSF, 1× Protease Inhibitor Cocktail (Merck Millipore 4693159001), and a phosphatase inhibitor cocktail containing 50 mM NaF, 2 mM Na_3_VO_4_, and 10 mM glycerolphosphate.

Phosphorylated BAF isoforms were resolved using Phos-tag SDS-PAGE as previously described (36) and analyzed by western blotting using rabbit polyclonal αBAF (1:2500). Protein band intensities from Phos-tag Western blot gels were quantified using a GS-800 Calibrated Densitometer and Quantity One 1-D analysis software (Bio-Rad).

### Immunostaining experiments

Immunostaining experiments were performed as described before (36). Briefly, cells were treated for 6 h with 25 μM colchicine (Sigma), immobilized onto a slide by centrifugation for 10 min at 500 rpm with low acceleration in a TermoShandon Cytospin using a single-chamber Cytofunnel and, then, fixed in 4% paraformaldehyde. Samples were immunostained with αBAF (1:300), rat αCENP-C (1:500), rabbit αCENP-C (1:300), αFlfl (1:1000), αWDB (1:2000), αMTS (1:500), and αTWS (1:100) antibodies. For visualization, slides were mounted in Mowiol (Merck Millipore 475904-M) containing 0.2 ng/ml DAPI (Merck Millipore D9542) and analyzed in a Leica TCS/SPE confocal microscope equipped with LAS/AF software. Images were acquired and processed identically using ImageJ 2.16.0/1.54p (http://imagej.nih.gov/ij/). Mean gray intensities were calculated on thresholded DAPI regions of interest running Analyze particles plugin on the FeatureJ Laplacian (http://imagescience.org /meijering/ software/ featurej/). Distance between centromeres were measured on centroids of paired CENP-C signals using the ‘Straight Line’ tool in ImageJ.

### Live imaging experiments

For live imaging, cells were grown in 35mm μ−dishes (ibidi 81156) coated with Concanavalin A. Images were taken with 63x magnification 1.46 NA oil-immersion lens with appropriate digital zoom in a Zeiss confocal LSM880 microscope equipped with Airyscan for image acquisition. Cells expressing GFP-and mCherry-tagged proteins were imaged sequentially (i.e., mCherry channel was acquired first, followed by GFP). Images were acquired every 120 seconds over a total period of 20 minutes. Each timepoint consisted of 165 z-slices, with a frame acquisition time of 218 ms. FastAiryscan raw data were preprocessed with the automatic setting of Zen Black and time-lapses were processed with Imaris 9 software. For 3D protein *foci* segmentation and tracking, time-lapse images containing protein *foci* to be identified were first bleach corrected (exponential fit), slightly smoothed (Fiji Gaussian filter), followed by a thresholding (Fiji RenyiEntropy). For tracking results, TrackMate 7 Fiji tool was used on mask images with the LAP tracker. Tracks were filtered based on the total number of spots detected and the presence of non-paired spot tracks. 3D tracking trajectories were generated with Matlab. The temporal offset between channel acquisitions was expected to result in a spatial shift of no more than 3 pixels, and was taken into account during data interpretation. Scripts, example data and parameters used could be found on Zenodo (https://zenodo.org/records/15490539). Coordinated movement between protein trajectories was evaluated using the discrete Fréchet distance, representing the maximum separation between two curves at any given point along their paths. Values were subsequently normalized to the corresponding nuclear diameter to ensure comparability across cells.

### Proximity Ligation Assay (PLA)

PLA was performed by using Duolink^R^ *in situ* detection Red kit (Merck Millipore DUO92008). First, cells were induced with 0.5 mM CuSO_4_ for 20-24h and, 6h before fixation with 4%PFA, cells were incubated with 25 μM colchicine. After fixation and permeabilization with 0.5% Triton X-100, samples were incubated with Duolink blocking solution in a preheated humid chamber for 1h at 37°C. Following blocking, cells were incubated overnight at 4°C with primary antibodies diluted in Duolink Antibody Diluent. The following antibody pairs and dilutions were used: rabbit αCENP-C (1:500) with mouse αGFP (1:2000); rabbit αCENP-C (1:500) with mouse αFlag (1:2000); mouse αGFP (1:2000) with rabbit αBAF (1:300); and mouse αFlag (1:2000) with rabbit αBAF (1:300).

### Statistical analysis

For each experiment, the number of independent biological replicates, sample sizes and statistical test used are indicated in the corresponding figure legend.

## RESULTS

### PP2A is required for BAF dephosphorylation during interphase

During interphase, BAF undergoes cycles of phosphorylation and dephosphorylation that regulate, among other functions, the anchoring of heterochromatin to the NE (27). As a component of the NE, BAF binds heterochromatin and mediates its tethering to the NE. Phosphorylation disrupts this interaction, whereas dephosphorylation restores heterochromatin binding and NE anchoring.

To investigate the functional relevance of this cycle, we analyzed heterochromatin mobility in *Drosophila* S2 cells co-expressing mCherry::HP1a, as a heterochromatin marker, together with either YFP::BAF, which undergoes normal phosphorylation/dephosphorylation cycles (27), or the phospho-dead mutant YFP::BAF^T4A/S5A^ (27). 3D time-lapse tracking revealed a tight association between the trajectories of mCherry::HP1a and both YFP::BAF (**Figures 1A** and **1C**) and YFP::BAF^T4A/S5A^ (**Figures 1B** and **1C**) at heterochromatin. We next quantified mCherry::HP1a signal displacement from the initial position over time as a measure of overall heterochromatin mobility. Notably, expression of the phospho-dead YFP::BAF^T4A/S5A^ mutant markedly reduced heterochromatin mobility (**Figure 1D**). Consistent with this, previous work showed that preventing BAF phosphorylation induces heterochromatin fragmentation, reinforces NE anchoring and increases nuclear instability (27). Together, these findings highlight the importance of dynamic BAF phosphorylation and dephosphorylation during interphase.

**Figure 1.**
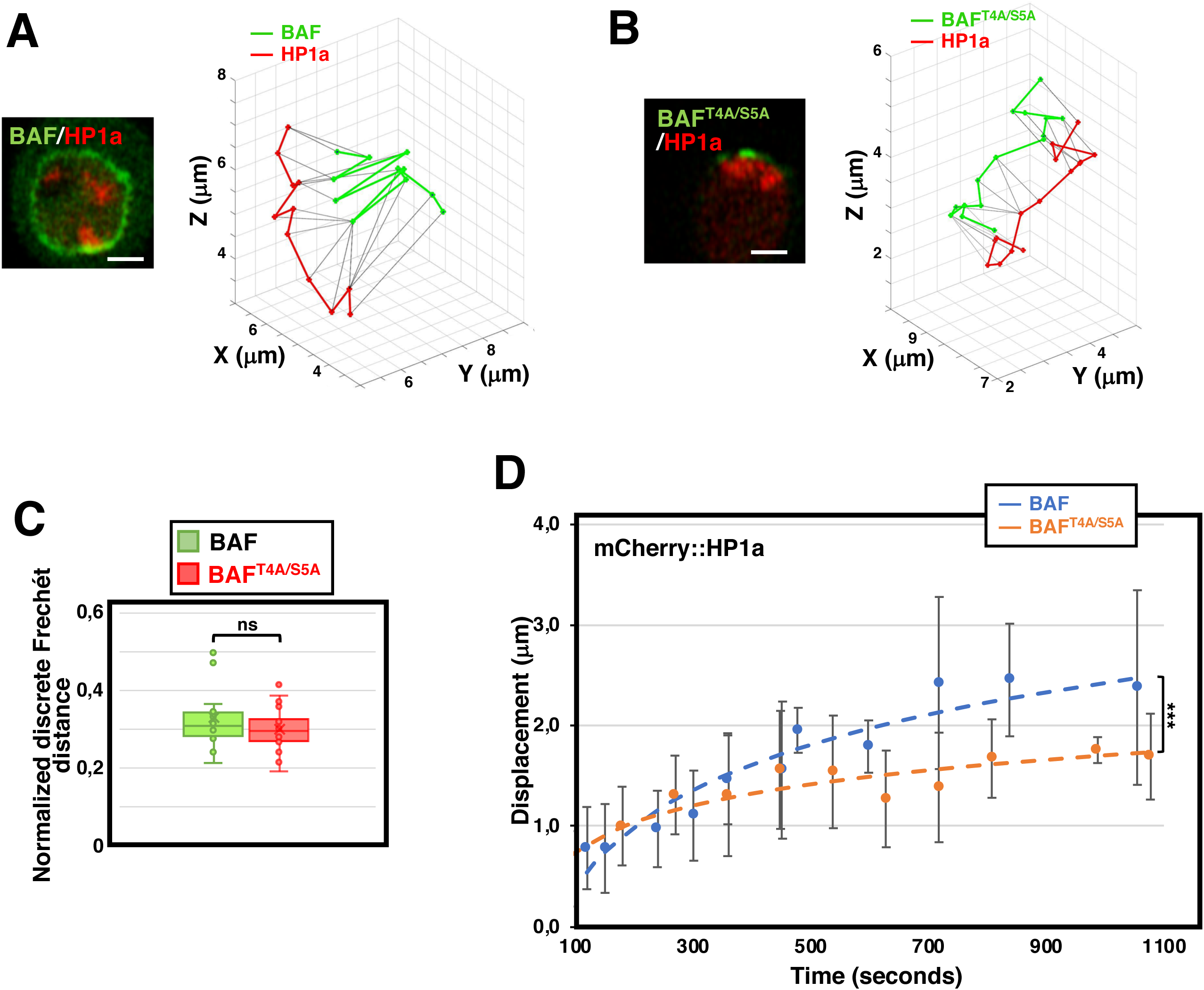
BAF phosphorylation controls mobility of heterochromatin. (**A-B**) Single time-lapse tracks for mCherry::HP1a (in red) and YFP::BAF (A) or YFP::BAF^T4A/S5A^ (B) (in green) at centromeric heterochromatin are presented. Experiments were performed in stably transfected S2 cell lines co-expressing mCherry::HP1a and either YFP::BAF (A) or YFP::BAF^T4A/S5A^ (B). In the left, the localization patterns of YFP::BAF and YFP::BAF^T4A/S5A^ are shown. Scale bars are 2 μm. Note that, as previously reported (27), while YFP::BAF is distributed around the entire NE, forming a perinuclear ring that includes the heterochromatin-anchoring domain, YFP::BAF^T4A/S5A^ is largely restricted to heterochromatin. (**C**) Box plots showing the degree of coordinated movement of mCherry::HP1a and either YFP::BAF or YFP::BAF^T4A/S5A^ quantified using the normalized discrete Fréchet distance, which calculates the upper bound of the largest distance between the corresponding polygonal trajectory curves. Boxes represent the median and interquartile range (IQR), whiskers are 1.5 * IQR (N = 18 trajectories from individual cells for each construct; p-value: ns > 0.05; unpaired Student’s t-test). (**D**) The displacement of the mCherry::HP1a signal from its initial position over a fixed time window are presented for cells expressing YFP::BAF or YFP::BAF^T4A/S5A^. Results are the mean of > 15 trajectories from individual cells for each construct. Error bars are SD. Differences in the mean distances across the groups were assessed using an Analysis of Covariance (ANCOVA). The covariate Time (X) is logarithmically transformed to fit the observed data pattern. Tukey’s Honestly Significant Difference (HSD) post-hoc test was used for pairwise comparisons (p-value: *** < 0.001).

It is well established that BAF phosphorylation is mediated by the VRK1/NHK1 kinase (28,31,41). In contrast, although earlier studies have shown that both PP2A and PP4 can dephosphorylate BAF (34,38,42), their specific contributions to this process remain poorly understood. To address this question, we induced BAF hyperphosphorylation in *Drosophila* S2 cells by treatment with okadaic acid (OA), a specific inhibitor of PP2A-like protein phosphatases (including PP2Ac and PP4c) resulting in mitotic arrest (51,52). Following OA washout, we monitored the recovery of the steady-state phosphorylation pattern as cells re-entered interphase. These experiments were performed in control mock-depleted cells and in cells depleted of the catalytic subunits of either PP2A (MTS) or PP4 (PP4c). The extent of BAF phosphorylation was determined by Phos-tag gel electrophoresis (53), which resolved mono- (1P) and di-phosphorylated (2P) BAF species from the non-phosphorylated (noP) form (27,36).

In control mock-depleted cells, BAF phosphorylation returned to basal levels within 6 hours after OA removal (**Figures 2A** and **2B**). A similar recovery was observed in PP4c-depleted cells (**Figures 2A** and **2B**). In contrast, MTS depletion significantly delayed the recovery of the basal phosphorylation pattern (**Figures 2A** and **2B**), as did depletion of the PP2A regulatory B/B’ subunits WDB and TWS (**Figures 2C** and **2D**). Altogether, these results identify PP2A^WDB^ and PP2A^TWS^ complexes as the main phosphatases responsible for BAF dephosphorylation during interphase, whereas PP4 plays only a minor role.

**Figure 2.**
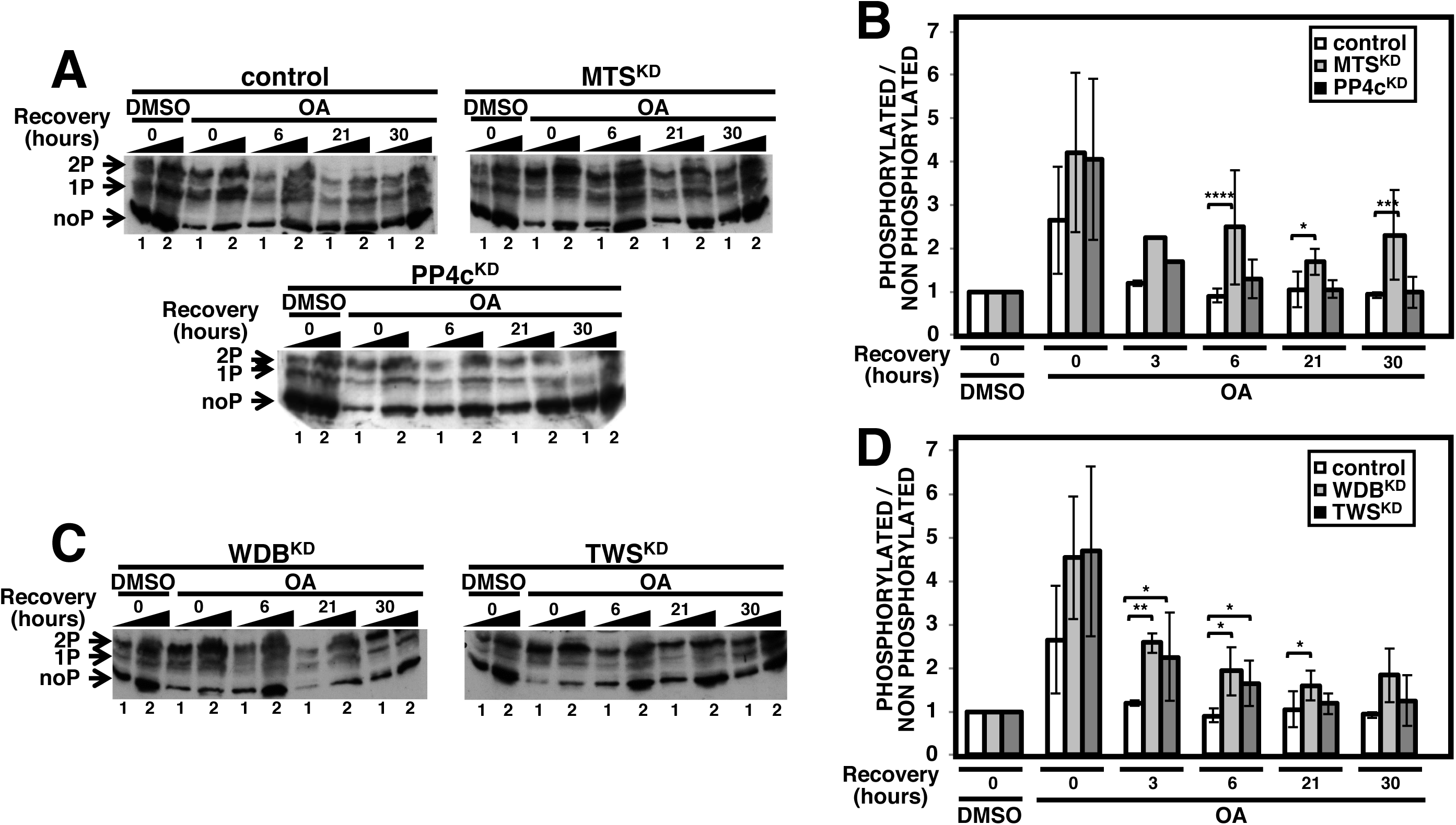
PP2A is required for BAF dephosphorylation during interphase. (**A**) The patterns of phosphorylation of BAF after treatment with okadaic acid (OA) are analyzed by Phos-tag gel electrophoresis of increasing amounts of extracts (lanes 1 and 2) prepared from control mock depleted cells (top left) and MTS (top right) or PP4c (bottom) knockdown cells. Cells were treated for 3h with either DMSO or 20nM OA, and harvested at increasing recovery times after washout (0h, 6h, 21h and 30h). Extracts were analyzed by WB using αBAF antibodies. The positions corresponding to non-phosphorylated (noP), and mono- (1P) and di-phosphorylated (2P) species are indicated. (**B**) Quantification of the results shown in A. The ratio of phosphorylated *versus* non phosphorylated forms is presented. Results are the average of 3-4 independent experiments. Errors bars are SD. (p-value: * < 0.05, *** < 0.001,**** < 0.0001; two-tailed paired Student’s t-test). (**C**) As in A, but for WDB (left) and TWS (right) knockdown cells. (**D**) Quantification of the results shown in C. The ratio of phosphorylated *versus* non phosphorylated forms is presented. Results are the average of 3-4 independent experiments. Errors bars are SD. (p-value: * < 0.05, ** < 0.01; two-tailed paired Student’s t-test).

### PP2A is required for centromeric localization of PP4, which directly interacts with centromeric BAF during mitosis

During mitosis, BAF becomes extensively phosphorylated and is largely released from chromatin (29,31,33,34). However, a small fraction of BAF (cenBAF) apparently escapes phosphorylation and remains bound to centromeres (36). Notably, both Flfl and WDB have been shown to localize at centromeres during mitosis (36,50,54,55), raising the possibility that PP2A^WDB^ and/or PP4^Flfl^ actively maintain cenBAF in an unphosphorylated state and, hence, bound to chromatin.

Immunostaining experiments confirmed the centromeric localization of both Flfl and WDB during mitosis (**Supplementary Figures S3A** and **S3B**), whereas TWS was not detectable at centromeres (**Supplementary Figure S3C**). However, while Flfl strongly overlapped with CENP-C (**Supplementary Figure S3A**, enlarged image), WDB was positioned adjacent to CENP-C, primarily in the interchromatid region (**Supplementary Figure S3B**, enlarged image). Consistently, proximity ligation assays (PLA) showed that Flfl, but not WDB, directly interacts with cenBAF. For these experiments, we used stable S2 cell lines co-expressing GFP::CENP-C, which localizes to centromeres and enables their direct visualization, and either Flag-tagged Flfl, WDB, or TWS. We then quantified the percentage of PLA-positive centromeres using αBAF and αFlag antibodies. In Flag::Flfl-expressing cells, ∼28% of centromeres displayed an αFlag/αBAF PLA signal (**Figures 3A**, top panel, and **3B**). This proportion was comparable to that observed in positive-control PLA experiments performed with αGFP, to detect GFP::CENP-C, and either αBAF (∼30%) (**Figure 3B** and **Supplementary Figure S4A**, top) or αFlag (∼38%) (**Figure 3B** and **Supplementary Figure S4A**, center), although lower than that observed in assays using αGFP and αCENP-C antibodies targeting the same GFP::CENP-C molecule (**Figure 3B** and **Supplementary Figure S4A**, bottom). In contrast, αFlag/αBAF PLA-positive centromeres in Flag::WDB- and Flag::TWS-expressing cells were markedly reduced (**Figures 3A**, center and bottom, and **3B**) and close to negative control levels observed in cells lacking any Flag-tagged construct (**Supplementary Figure S4B**). Altogether, these results indicate that PP4^Flfl^, but not PP2A^WDB^, directly interacts with cenBAF and may therefore actively counteract its phosphorylation.

**Figure 3.**
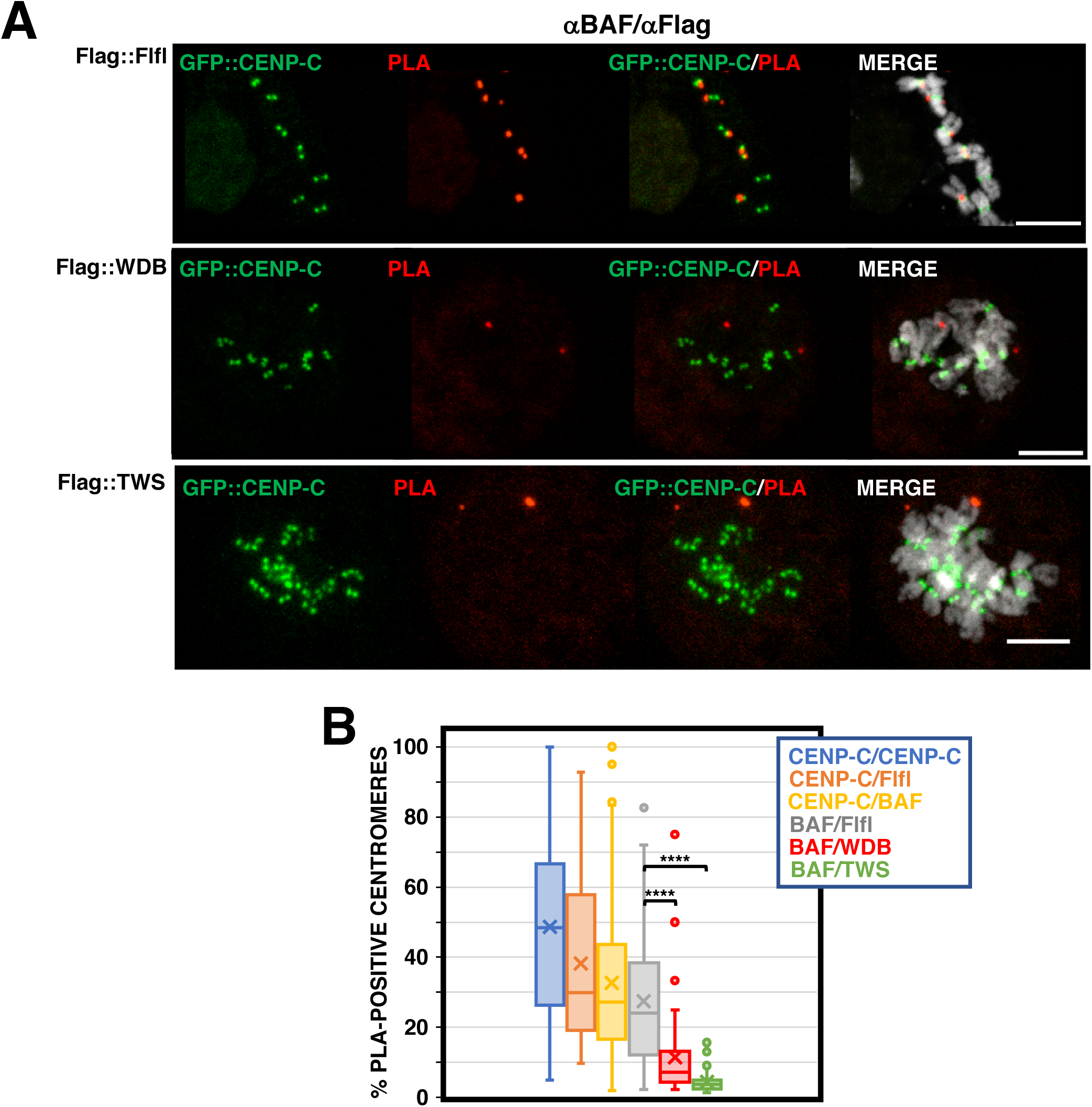
Flfl interacts with centromeric BAF (cenBAF). (**A**) PLA experiments performed with αFlag and αBAF antibodies in stable S2 cell lines co-expressing eGFP::CENP-C (direct fluorescence in green) and either 3xFlag::Flfl (top), 3xFlag::WDB (center) or 3xFlag::TWS (bottom). Specific PLA signals, indicating close proximity between the corresponding proteins, are shown in red. DNA was stained with DAPI (in grey). Scale bars are 5 μm. (**B**) Quantification of the results shown in A. Box plots showing the percentage of centromeres displaying αBAF/αFlag signal in cells expressing 3xFlag::Flfl (BAF/Flfl, grey), 3xFlag::WDB (BAF/WDB, red) or 3xFlag::TWS (BAF/TWS, green). The results from control positive PLA experiments are included for comparison: CENP-C/CENP-C (blue; see Supplementary Figure S4A, bottom), CENP-C/Flfl (brown; see Supplementary Figure S4A, center) and CENP-C/BAF (yellow; see Supplementary Figure S4A, top). Boxes represent the median and interquartile range (IQR), whiskers are 1.5 * IQR. Results are from 4 independent experiments (N > 50 for each condition; p-values: **** < 0.0001; two-tailed paired Student’s t-test).

Notably, WDB is required for centromeric localization of Flfl, as depletion of WDB strongly reduced centromeric Flfl levels in comparison to control cells (**Figures 4A** and **4B**). A similar reduction was observed upon depletion of MTS (**Figure 4B** and **Supplementary Figure S5A**), whereas TWS depletion did not reduce centromeric Flfl levels (**Figure 4B** and **Supplementary Figure S5B**). Importantly, centromeric Flfl levels in WDB-depleted cells were comparable to those observed following direct Flfl depletion (**Figure 4B** and **Supplementary Figure S5C**), while total Flfl levels remain unchanged (**Supplementary Figures S2E** and **S2F**).

**Figure 4.**
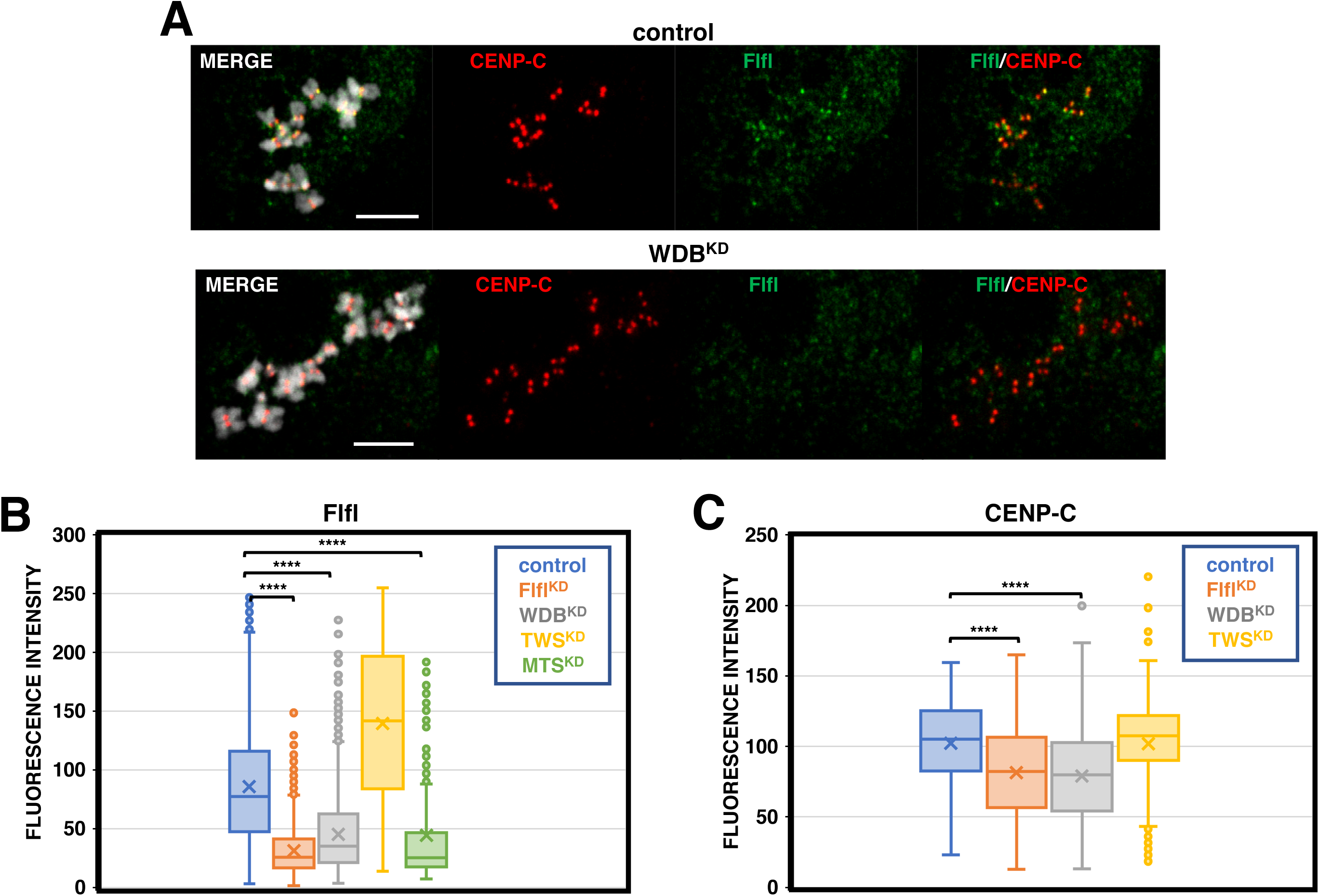
Centromeric localization of Flfl depends on WDB. (**A**) Immunostainings of metaphase chromosomes with αCENP-C (in red) and αFlfl (in green) performed in control mock depleted (top) and WDB knockdown (bottom) cells. DNA was stained with DAPI (in grey). Scale bars are 5 μm. (**B**) Quantification of the results shown in A. Box plots showing the mean αFlfl fluorescence intensity at the centromere in control mock depleted cells (blue) and in WDB knockdown (grey) cells. The results obtained in cells depleted for Flfl (brown; see Supplementary Figure S5C), TWS (yellow; see Supplementary Figure S5B) and MTS (green; see Supplementary Figure S5A) are included. Boxes represent the median and interquartile range (IQR), whiskers are 1.5 * IQR. Results are from 3 independent experiments. (N > 1200 for each condition; p-values: **** < 0.0001; two-tailed paired Student’s t-test). (**C**) As in B, but for mean αCENP-C fluorescence intensity in mock depleted cells (blue; see Supplementary Figure S6A) and in cells depleted for Flfl (brown; see Supplementary Figure S6B), WDB (grey; see Supplementary Figure S6C) or TWS (yellow; see Supplementary Figure S6D). Boxes represent the median and interquartile range (IQR), whiskers are 1.5 * IQR. Results are from 3 independent experiments. (N > 1200 for each condition; p-values: **** < 0.0001; two-tailed paired Student’s t-test).

Given that Flfl stabilizes CENP-C at centromeres (36), we next examined whether WDB depletion affects centromeric CENP-C levels and, indeed, we found a comparable reduction as upon direct Flfl depletion (**Figure 4C** and **Supplementary Figures S6A-S6C**). In contrast, depletion of TWS did not reduce centromeric CENP-C levels (**Figure 4C** and **Supplementary Figures S6D**). Interestingly, although only slightly, Flfl depletion also reduced centromeric WDB levels (**Supplementary Figure S7**).

### PP4 depletion induces premature BAF re-association with chromosomes

During mitotic exit, the large pool of phosphorylated BAF (pBAF) is dephosphorylated and re-associates with chromatin, forming a perichromosomal layer that surrounds the bulk of chromosomes (14,29,34,36–40). At metaphase, BAF is largely restricted to centromeres (36) (**Figure 5A**, top). However, we observed that approximately 25% of chromosomes already displayed a perichromosomal BAF layer at this stage (**Figure 5B**). Analysis of chromosome frequency distribution as a function of interchromatid distance at the centromeres revealed a shift toward larger distances in chromosomes with perichromosomal BAF compared with those lacking it (**Figure 5C**, top), suggesting that BAF re-association begins in early anaphase, as chromosomes start to segregate.

**Figure 5.**
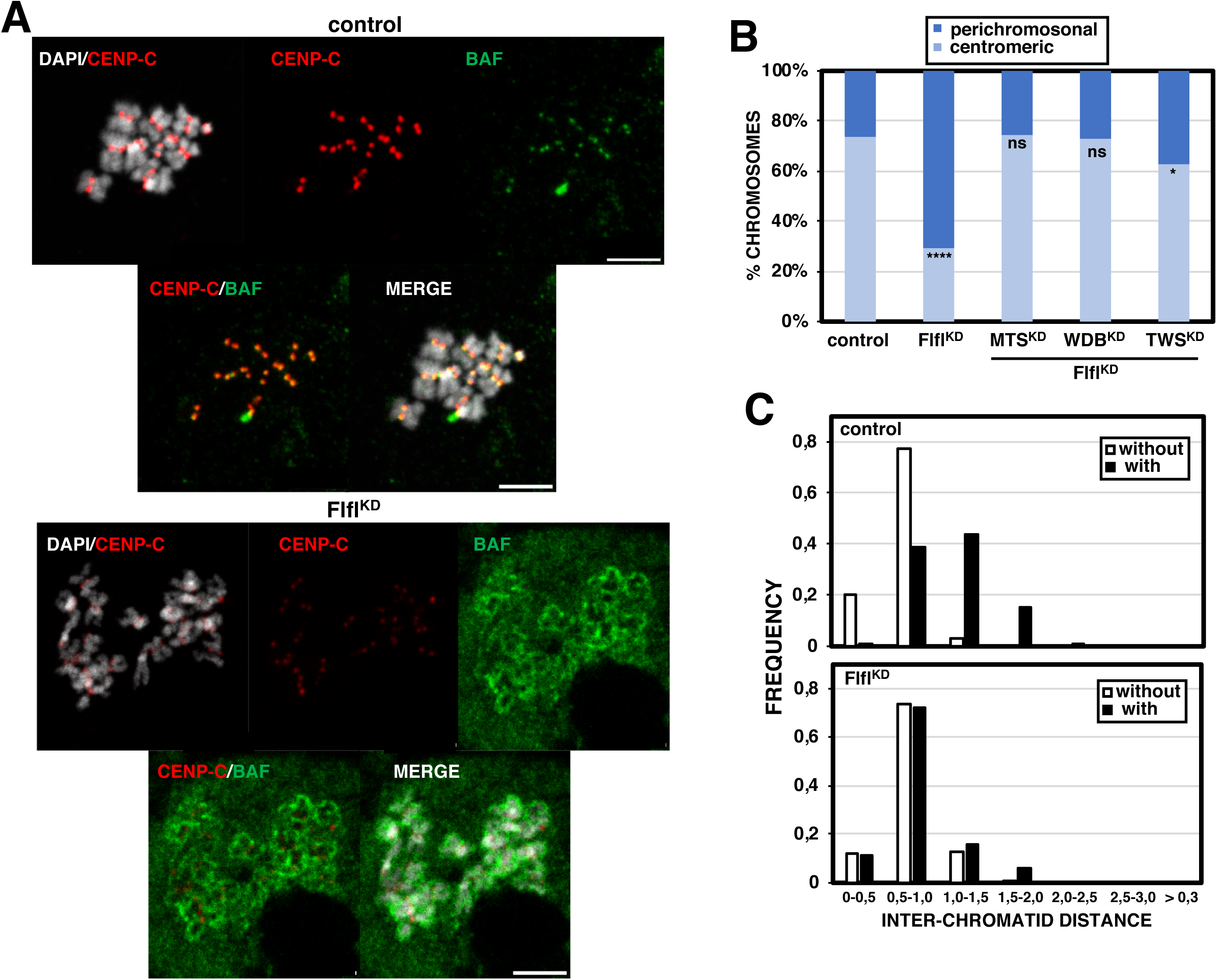
Flfl depletion induces premature BAF re-association with chromosomes. (**A**) Immunostainings of metaphase chromosomes with αCENP-C (in red) and αBAF (in green) antibodies performed in control mock depleted (top) and Flfl-depleted cells (bottom). DNA was stained with DAPI (in grey). Scale bars are 5 μm. (**B**) Quantification of the results shown in A. The percentages of chromosomes showing a layer of perichromosomal BAF surrounding the chromosomes and those where BAF is detected only at centromeres are shown for control and Flfl-depleted cells. Results from double knockdown cells for Flfl and either MTS, WDB or TWS are also included. Results are the average of 5-8 independent experiments (N > 180 for each condition; p-values respect to control: ns > 0.05; * < 0.05; **** < 0.0001; Fisher’s exact test) (**C**) The chromosome frequency distribution as a function of interchromatid distance at the centromeres is presented for chromosomes showing perichromosomal BAF and not in control mock depleted (top) and Flfl-depleted cells (bottom). Results are from 3-6 independent experiments. (N > 600 for each condition).

Notably, Flfl depletion strongly increased the proportion of chromosomes displaying perichromosomal BAF (36) (**Figures 5A**, bottom, and **5B**), which, unlike in control cells, exhibited a frequency distribution as a function of interchromatid distance indistinguishable from that of chromosomes lacking perichromosomal BAF (**Figure 5C**, bottom). These results indicate that Flfl depletion triggers premature BAF re-association at or before the metaphase-to-anaphase transition.

Interestingly, depletion of either MTS or WDB rescued the increase in perichromosomal BAF observed upon Flfl depletion (36) (**Figure 5B**), indicating that PP2A^WDB^ is required for dephosphorylation of mitotic pBAF and the subsequent formation of the perichromosomal BAF layer. Consistently, WDB depletion alone reduced the proportion of chromosomes displaying perichromosomal BAF (**Figures 6A**, top, and **6B**). In addition, WDB depletion induced premature BAF re-association with chromosomes (**Figure 6C**, center), resembling the phenotype observed upon Flfl depletion (**Figure 5C**, bottom). This effect likely reflects the requirement of WDB for centromeric Flfl localization (see **Figure 4**). Together, these findings reveal a dual role for PP2A^WDB^ in BAF re-association: it promotes pBAF dephosphorylation and re-association with chromosomes, while also controling the timing of this process through the regulation of centromeric PP4^Flfl^ localization.

**Figure 6.**
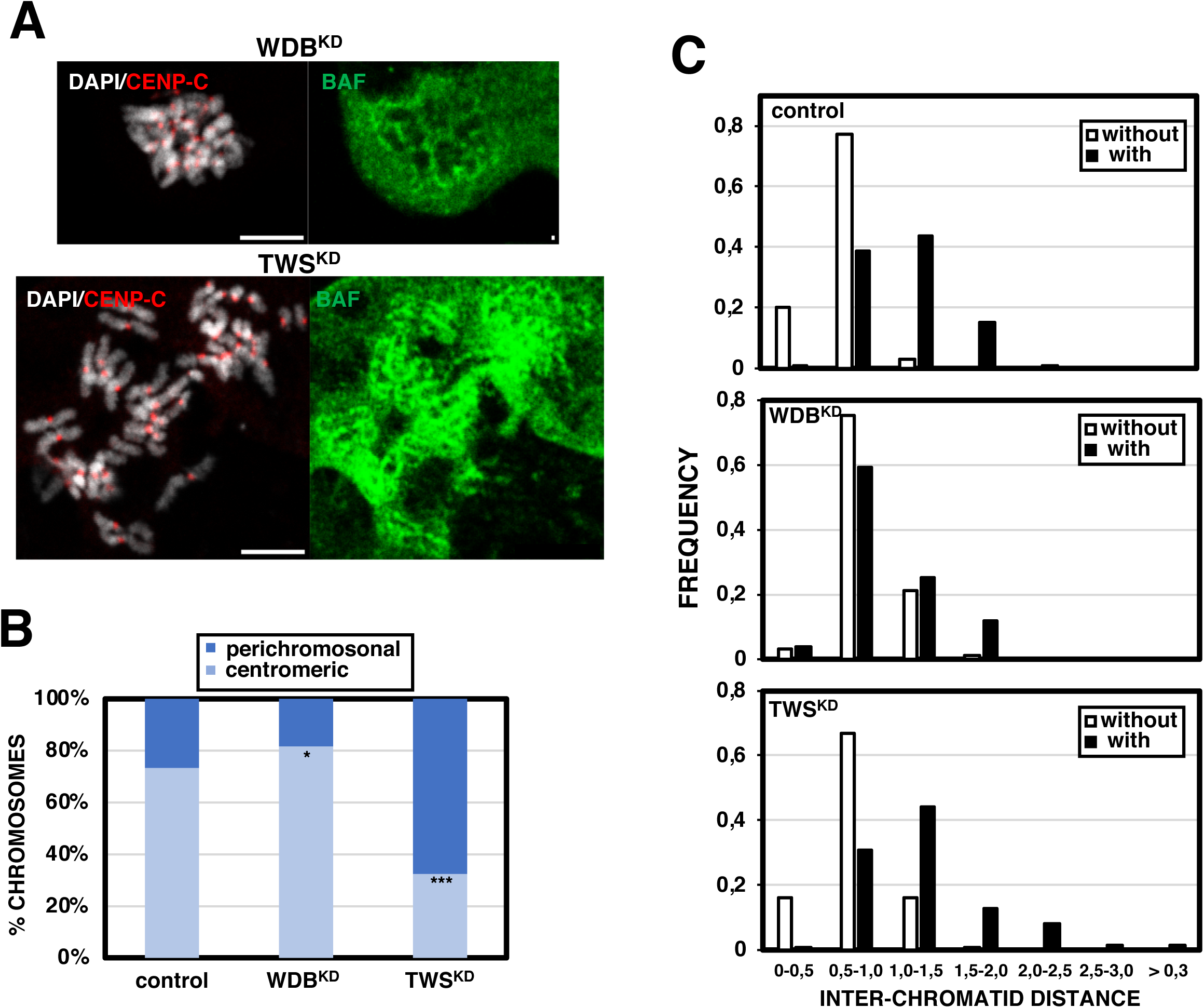
Effects of PP2A depletion on BAF re-association with chromosomes. (**A**) Immunostainings of metaphase chromosomes with αCENP-C (in red) and αBAF (in green) antibodies performed in WDB-depleted (top) and TWS-depleted cells (bottom). DNA was stained with DAPI (in grey). Scale bars are 5 μm. (**B**) Quantification of the results shown in A. The percentages of chromosomes showing a layer of perichromosomal BAF surrounding the chromosomes and those where BAF is detected only at centromeres are shown for control, WDB-depleted and TWS-depleted cells. Results are the average of 4-8 independent experiments. (N > 180 for each condition; p-values respect to control: * < 0.05, *** < 0.001; Fisher’s exact test). (**C**) The chromosome frequency distribution as a function of interchromatid distance at the centromeres is presented for chromosomes showing perichromosomal BAF and not in control (top), WDB-depleted (center) and TWS-depleted (bottom) cells. Results are from 3-6 independent experiments. (N > 560 for each condition).

In contrast, TWS depletion produced markedly different effects. Although it strongly increased the proportion of chromosomes displaying perichromosomal BAF (**Figures 6A**, bottom, and **6B**), it did not alter the timing of BAF re-association (**Figure 6C**, bottom). Moreover, compared with depletion of either MTS or WDB, TWS depletion only partially rescued the increase in perichromosomal BAF observed in Flfl-depleted cells (**Figure 5B**). Together, these findings suggest that TWS makes only a minor contribution to pBAF dephosphorylation at this stage. Rather, they are consistent with the proposed role of TWS in completion of mitotic exit (56,57), as TWS depletion delays progression through anaphase and consequently leads to the accumulation of chromosomes at these stages (58).

## DISCUSSION

Previous studies have shown that both PP2A and PP4 dephosphorylate BAF (34,38,42), but their specific contributions to BAF dephosphorylation remain poorly understood. Here, we show that PP2A is the major protein phosphatase involved in BAF dephosphorylation, whereas PP4 likely acts during mitosis.

Our results show that recovery of the normal pattern of BAF phosphorylation after OA treatment, which arrests cells in mitosis (51,52), is strongly delayed in cells depleted of either the PP2A catalytic subunit MTS or the regulatory B/B’ subunits WDB and TWS. These findings indicate that, during progression into interphase, both PP2A^WDB^ and PP2A^TWS^ holoenzymes are involved in BAF dephosphorylation. Whether both complexes are active in BAF dephosphorylation throughout interphase or instead act at specific stages or locations remains to be determined. In this regard, it has been reported that PP2A^TWS^ is specifically involved in BAF dephosphorylation in late anaphase (38).

PP2A-mediated BAF dephosphorylation likely regulates the anchoring of heterochromatin to the NE in interphase. Heterochromatin anchoring is a highly dynamic process that requires rapid cycles of BAF phosphorylation and dephosphorylation (27). Impairing BAF phosphorylation, reinforces anchoring and, as shown in this study, strongly reduces heterochromatin mobility, which results in the accumulation of multiple defects and nuclear instability (27). Further work is required to dissect the role of PP2A in regulating chromatin-NE interactions.

Restoring normal levels of BAF phosphorylation after OA treatment is not significantly affected by PP4 depletion, suggesting that this protein phosphatase makes only a minor contribution to BAF dephosphorylation during interphase. The contribution of PP4 appears to be largely restricted to mitosis, where the PP4 regulatory 3 subunit Flfl associates with centromeres in a CENP-C-dependent manner (50). Consistent with this, we show here that Flfl directly interacts *in vivo* with the small pool of cenBAF that remains bound to centromeres during mitosis, suggesting that PP4 is likely involved in maintaining cenBAF unphosphorylated and associated with centromeric chromatin. Accordingly, Flfl has been shown to be required for stabilization of cenBAF at centromeres (36).

Notably, as shown here and elsewhere (54,55), WDB also localizes to centromeric regions on metaphase chromosomes. However, it preferentially accumulates at the interchromatid region and PLA experiments failed to detect a direct interaction with cenBAF, thereby excluding a potential role of PP2A^WDB^ in maintaining cenBAF unphosphorylated. Importantly, centromeric localization of Flfl requires WDB, although the mechanism underlying this effect remains unknown. Flfl contains a putative WDB-binding short linear motif (aa 143-151: LEDISETIQ), raising the possibility of a direct WDB-Flfl interaction. Further work will be required to clarify the molecular basis of this regulation.

During mitotic exit, PP2A^WDB^ and PP2A^TWS^ act in coordinated phosphatase programs, with PP2A^TWS^ executing later dephosphorylation events required for completion of mitosis (56,57). In this context, it has been shown that PP2A^TWS^ drives BAF dephosphorylation at late anaphase (38), a process that constitutes an essential step for NER (14,29,34,37–40). However, upon TWS depletion, BAF dephosphorylation and NER are delayed rather than completely abolished (38), suggesting that additional phosphatase complexes contribute to BAF dephosphorylation during mitotic exit. Notably, as shown here and previously reported (36), a perichromosomal layer of BAF is already detectable at very early anaphase, coincident with the onset of chromosome segregation. At this stage, PP2A^TWS^ is thought to be largely inactive. Consistently, we found that TWS depletion does not impair formation of the early perichromosomal BAF layer. In contrast, WDB depletion strongly disrupts this process, supporting the idea that the initial BAF dephosphorylation events occurring during early anaphase are mediated primarily by PP2A^WDB^. Importantly, the timing of this process depends on PP4^Flfl^, as Flfl depletion leads to premature BAF re-association at, or even before, transition to anaphase. Altogether, these observations suggest a finely tune interplay between PP4 and PP2A in regulating BAF dephosphorylation and chromatin re-association during mitotic exit. Further work will be required to dissect this regulatory interplay in detail.

## ACKNOWLEDGEMENTS

We are thankful to Dr. Vincent Archambault and Dr. Amita Sehgal for αTWS and αWDB antibodies, respectively, and to the *Drosophila* Genomics Resource Center (supported by NIH grant no. 2P40OD010949) for the cDNA clones and *Drosophila* Gateway plasmids. This work was financed by grants PGC2018-094538-B-100 and PID2021-123303NB-I00 from MICIN/AEI 10.13039/501100011033 and “FEDER, una manera de hacer Europa”, and of the Generalitat de Catalunya (SGR2017-475) to F.A., and Momentum Program Grant (LP2017-7/2017) from the Hungarian Academy of Sciences to Z.L.. P.E-F. acknowledges receipt of an FPI fellowship from MICIN/AEI 10.13039/501100011033 and “FEDER, una manera de hacer Europa”.

## DATA AVAILABILITY

The data underlying this article will be shared upon reasonable request to the corresponding authors.

## AUTHORS CONTRIBUTIONS

Conceptualization, M.T-L. and F.A.; Investigation, M.T-L., P.E-F., C.Z.; Resources, E.A. and Z.L.; Writing-Original Draft, M.T-L. and F.A; Writing-Review & Editing, E.A. and Z.L; Supervision, M.T-L. and F.A.; Funding Acquisition F.A. and Z.L.

## Notes

### Competing Interest Statement

The authors have declared no competing interest.

