## Supplemental for "Coordinated regulation of Barrier-to-Autointegration Factor (BAF) phosphorylation by protein phosphatases PP2A and PP4"

**SUPPLEMENTARY INFORMATION**

**Supplementary Table S1.** Primers used for dsRNAs synthesis in RNAi knockdown experiments and for plasmid generation.

| | |
|---|---|
| dsLacZ fw | TAATACGACTCACTATAGGGATGACCATGATTACGCCAAGC |
| dsLacZ rev | TAATACGACTCACTATAGGGCAATTTCCATTCGCCATTCAG |
| dsFlfI fw | TAATACGACTCACTATAGGGAGAATGACGACTGACACCCGC |
| dsFlfI rev | TAATACGACTCACTATAGGGAGAACAACTTTTCCTTTCGCA |
| dsMTS fw | TAATACGACTCACTATAGGGGCCAAGGAGATTCTCTCC |
| dsMTS rev | TAATACGACTCACTATAGGGCTGGCCAAAGGTGTAACC |
| dsPP4c fw | TAATACGACTCACTATAGGGAGAATGTCCGACTACAGCGAC |
| dsPP4c rev | TAATACGACTCACTATAGGGAGACCACACGGCCGTCGATCC |
| dsTWS fw | TAATACGACTCACTATAGGGAGACTGATCCGGGATCCACAGAATGTAA |
| dsTWS rev | TAATACGACTCACTATAGGGAGACACACTTTGATGCTCAAGTAATCCC |
| dsWDB1 fw | TAATACGACTCACTATAGGGAGATCGATCCGCCGCAGTTTGTCAAGAT |
| dsWDB1 rev | TAATACGACTCACTATAGGGAGATCGACTGCTGCTGATGAGAGTTCAG |
| dsWDB2 fw | TAATACGACTCACTATAGGGAGAGCTCAAGAAGAAGGGTAAAAAGAGT |
| dsWDB2 rev | TAATACGACTCACTATAGGGAGACCATATATACGATGCAATACCGTCT |
| dsWDB2 rev | TAATACGACTCACTATAGGGAGACCATATATACGATGCAATACCGTCT |
| GW-FlfI-fw | GGGGACAAGTTTGTACAAAAAAGCAGGCTTAATGACGACTGACACCCGCCGACGCG |
| GW-FlfI-rev | GGGGACCACTTTGTACAAGAAAGCTGGGTACTATGCCTGACGCGCGCGCTTTTGT |
| mts_GW_fw | GGGGACAAGTTTGTACAAAAAAGCAGGCTTAATGGAGGATAAAGCAACAACAAAAG |
| mts_GW_Rev | GGGGACCACTTTGTACAAGAAAGCTGGGTATTAAAGGAAATAATCGGGTGTTCTT |
| Wdb_GW_fw | GGGGACAAGTTTGTACAAAAAAGCAGGCTTAATGTCATCGGGCACGTTTGTGGATC |
| Wdb_GW_Rev | GGGGACCACTTTGTACAAGAAAGCTGGGTATTAGTTGTCCGCCTTATCCTGTTTG |
| Tws-RA_GW_fw | GGGGACAAGTTTGTACAAAAAAGCAGGCTTAATGGGTCGCTGGGGACGGCAGAGTC |
| Tws-RA_GW_Rev | GGGGACCACTTTGTACAAGAAAGCTGGGTACTAAAATTTATCCTGAAATATGAAG |

### LEGENDS TO SUPPLEMENTARY FIGURES

**Supplementary Figure S1**. **Western blot analysis of co-expression of eGFP-CENP-C and 3×Flag-tagged proteins.** (**A-C**) Crude protein extract of double-stable D. Mel (2) cell lines co-expressing eGFP::CENP-C and either 3×Flag::WDB, or 3×Flag::TWS-RA (A), 3×Flag::MTS (B) or 3×Flag::Flfl (C) were fractionated by SDS-PAGE and analyzed by WB using $\alpha$GFP, $\alpha$CENP-C, or $\alpha$Flag antibodies as indicated. Control indicates the parental D. Mel (2) cell line. Uninduced (–) and 1 mM $CuSO_4$-induced (+) cells were compared. The position of Mw (kDa) markers is indicated.

**Supplementary Figure S2**. **Assessment of depletion efficiency in the RNAi knockdown experiments.** (**A**) WB analysis with $\alpha$MTS (top) and $\alpha$PP4c (bottom) antibodies of increasing amounts of total protein extracts (lanes 1 and 2) prepared from MTS (top) and PP4c (bottom) knockdown cells and from control mock depleted cells. $\alpha$Tubulin was used for loading control. The position of Mw (kDa) markers is indicated. (**B**) Quantification of the results shown in A. The levels of MTS and PP4c relative to Tubulin are shown. Results are the average of 3 independent experiments. Errors bars are SD. (p-value: ** $< 0.01$, **** $< 0.0001$; two-tailed paired Student's t-test). (**C**) WB analysis with $\alpha$WDB (top) and $\alpha$TWS (bottom) antibodies of increasing amounts of total protein extracts (lanes 1 and 2) prepared from WDB (top) and TWS (bottom) knockdown cells and from control mock depleted cells. $\alpha$Lamin was used for loading control. The position of Mw (kDa) markers is indicated. (**D**) Quantification of the results shown in C. The levels of WDB and TWS relative to Lamin are shown. Results are the average of 3 independent experiments. Errors bars are SD. (p-value: ** $< 0.01$, **** $< 0.0001$; two-tailed paired Student's t-test). (**E**) WB analysis with $\alpha$Flfl antibodies of increasing amounts of total protein extracts (lanes 1 and 2) prepared from control mock-depleted cells and cells depleted for Flfl (top, center), WDB (top, right) or TWS (bottom, right). $\alpha$H3 was used for loading control. The position of Mw (kDa) markers is indicated. (**F**) Quantification of the results shown in E. The levels of Flfl relative to H3 are shown. Results are the average of 3 independent experiments. Errors bars are SD. (p-values: ns > 0.05; **** < 0.0001; two-tailed paired Student's t-test).

**Supplementary Figure S3. Flfl and WDB localize to centromeres in metaphase chromosomes**. Immunostainings of metaphase chromosomes with αCENP-C (in red) and either αFlfl (**A**), αWDB (**B**) or αTWS (**C**) (in green) are presented. DNA was stained with DAPI (in grey). Scale bars are 5 μm. Enlarged images are shown on the right. Scale bars are 0.2 μm.

**Supplementary Figure S4. Control PLA experiments.** (**A**) Positive-control PLA experiments performed in cells expressing eGFP::CENP-C (direct fluorescence in green) and 3xFlag::Flfl with αGFP and either αBAF (top), αFlag (center) or αCENP-C (bottom) antibodies. The PLA signal is shown in red. DNA was stained with DAPI (in grey). Scale bars are 5 μm. (**B**) Negative-control PLA experiment performed with αGFP and αFlag antibodies in cells expressing eGFP::CENP-C (direct fluorescence in green) and no Flag-tagged construct. The PLA signal is shown in red. DNA was stained with DAPI (in grey). Scale bar is 5 μm.

**Supplementary Figure S5**. Immunostainings of metaphase chromosomes with αCENP-C (in red) and αFlfl (in green) performed in cells depleted for MTS (top), TWS (center) or Flfl (bottom). DNA was stained with DAPI (in grey). Scale bars are 5 μm.

**Supplementary Figure S6.** Immunostainings of metaphase chromosomes with αCENP-C antibodies (in red) performed in control mock depleted cells (A) and in cells depleted for Flfl (B), WDB (C) or TWS (D). DNA was stained with DAPI (in grey). Scale bars are 5 μm.

**Supplementary Figure S7**. (**A**) Immunostainings of metaphase chromosomes with αCENP-C (in red) and αWDB (in green) antibodies performed in control mock depleted cells (top) and Flfl depleted cells (bottom). DNA was stained with DAPI (in grey). Scale bars are 2 μm. (**B**) Quantification of the results shown in A.

Box plots showing the mean $\alpha$WDB fluorescence intensity at the centromere in control mock depleted cells (blue) and Flfl depleted cells (brown). Boxes represent the median and interquartile range (IQR), whiskers are 1.5 * IQR. Results are from 2 independent experiments. (N = 238 and 1342 for control and $Flfl^{KD}$, respectively; p-value: **** < 0.0001).

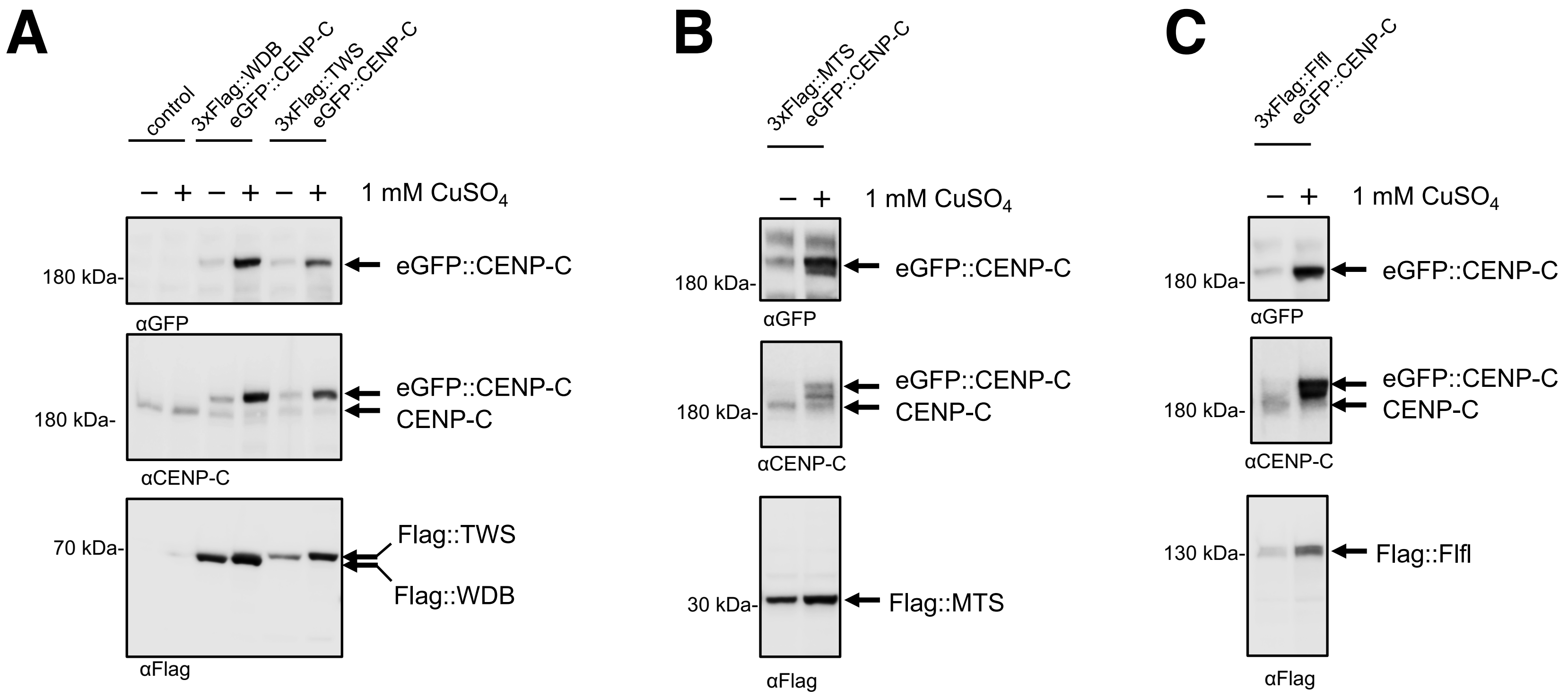
A
control
3xFlag::WDB eGFP::CENP-C
3xFlag::TWS eGFP::CENP-C
– + – + – + 1 mM $CuSO_4$
180 kDa-
eGFP::CENP-C
αGFP
180 kDa-
eGFP::CENP-C
CENP-C
αCENP-C
70 kDa-
Flag::TWS
Flag::WDB
αFlag
B
3xFlag::MTS eGFP::CENP-C
– + 1 mM $CuSO_4$
180 kDa-
eGFP::CENP-C
αGFP
180 kDa-
eGFP::CENP-C
CENP-C
αCENP-C
30 kDa-
Flag::MTS
αFlag
C
3xFlag::Flfl eGFP::CENP-C
– + 1 mM $CuSO_4$
180 kDa-
eGFP::CENP-C
αGFP
180 kDa-
eGFP::CENP-C
CENP-C
αCENP-C
130 kDa-
Flag::Flfl
αFlag

**Figure S1**

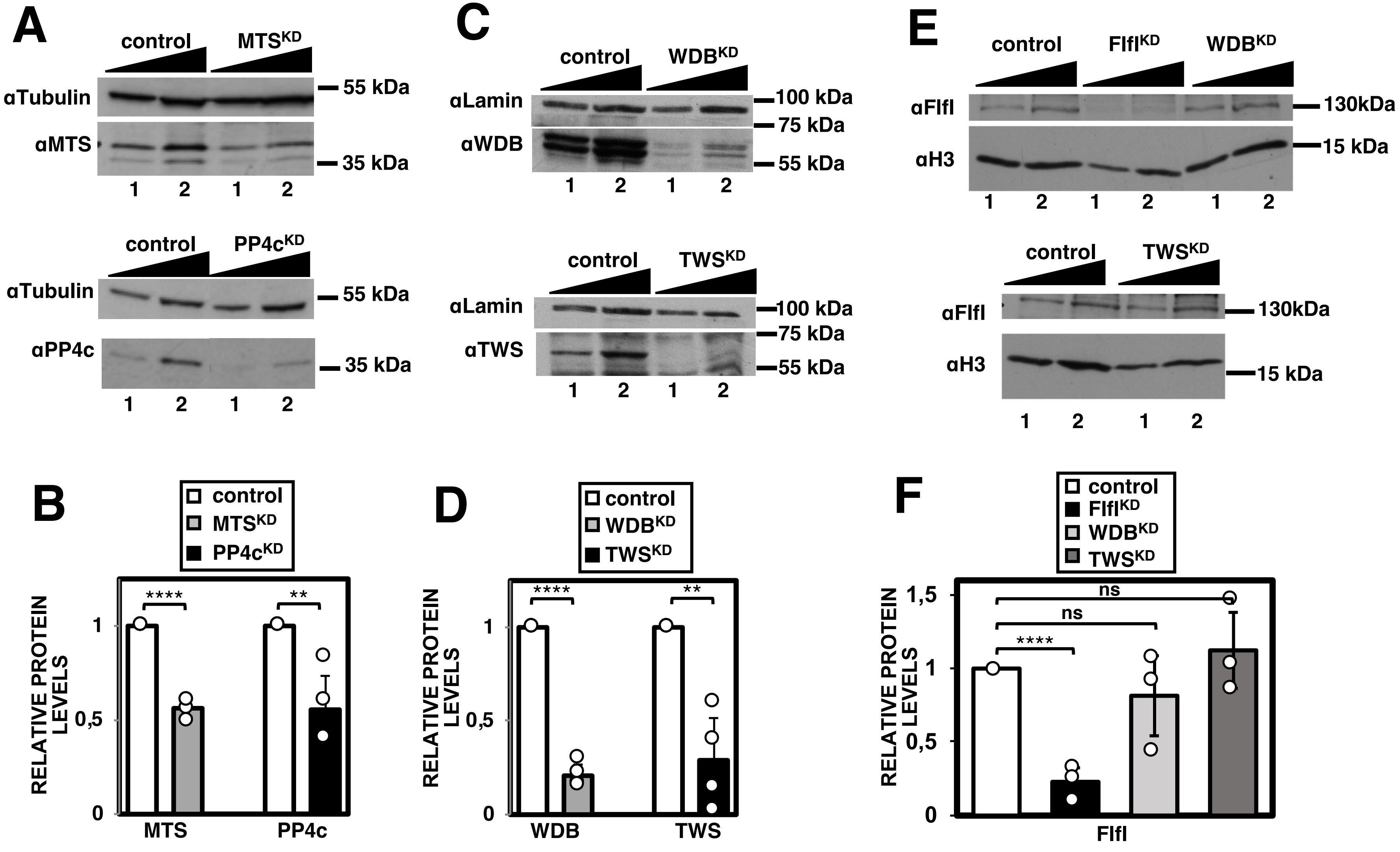
A
control
MTS^KD
αTubulin
55 kDa
αMTS
35 kDa
1 2 1 2
control
PP4c^KD
αTubulin
55 kDa
αPP4c
35 kDa
1 2 1 2
C
control
WDB^KD
αLamin
100 kDa
75 kDa
αWDB
55 kDa
1 2 1 2
control
TWS^KD
αLamin
100 kDa
75 kDa
αTWS
55 kDa
1 2 1 2
E
control
Flfl^KD
WDB^KD
αFlfl
130kDa
αH3
15 kDa
1 2 1 2 1 2
control
TWS^KD
αFlfl
130kDa
αH3
15 kDa
1 2 1 2
B
control
MTS^KD
PP4c^KD
RELATIVE PROTEIN LEVELS
****
**
1
0,5
0
MTS
PP4c
D
control
WDB^KD
TWS^KD
RELATIVE PROTEIN LEVELS
****
**
1
0,5
0
WDB
TWS
F
control
Flfl^KD
WDB^KD
TWS^KD
RELATIVE PROTEIN LEVELS
ns
ns
****
1,5
1
0,5
0
Flfl

Figure S2

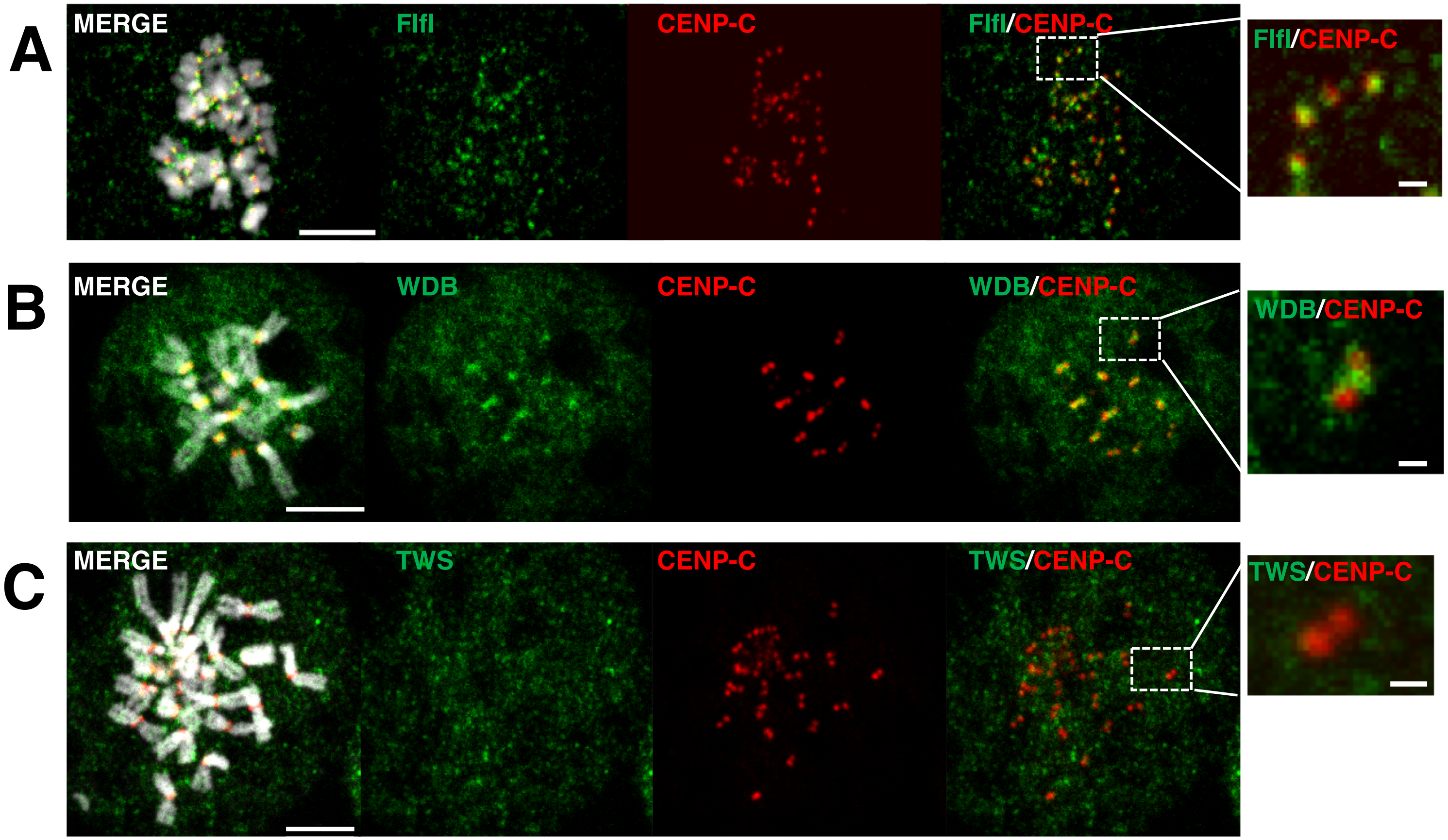
A
MERGE
Flfl
CENP-C
Flfl/CENP-C
Flfl/CENP-C
B
MERGE
WDB
CENP-C
WDB/CENP-C
WDB/CENP-C
C
MERGE
TWS
CENP-C
TWS/CENP-C
TWS/CENP-C

Figure S3

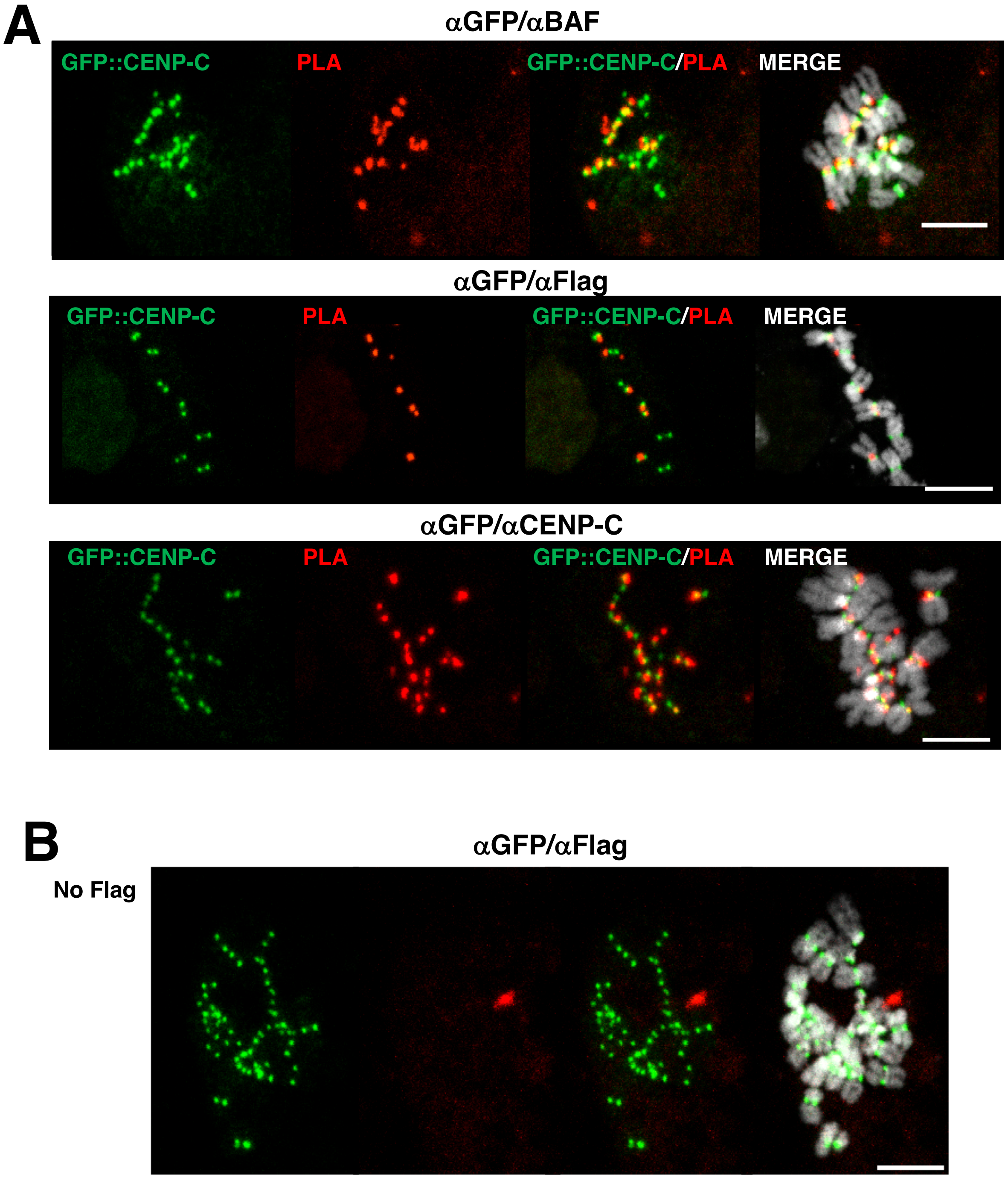
A
αGFP/αBAF
GFP::CENP-C
PLA
GFP::CENP-C/PLA
MERGE
αGFP/αFlag
GFP::CENP-C
PLA
GFP::CENP-C/PLA
MERGE
αGFP/αCENP-C
GFP::CENP-C
PLA
GFP::CENP-C/PLA
MERGE
B
αGFP/αFlag
No Flag

Figure S4

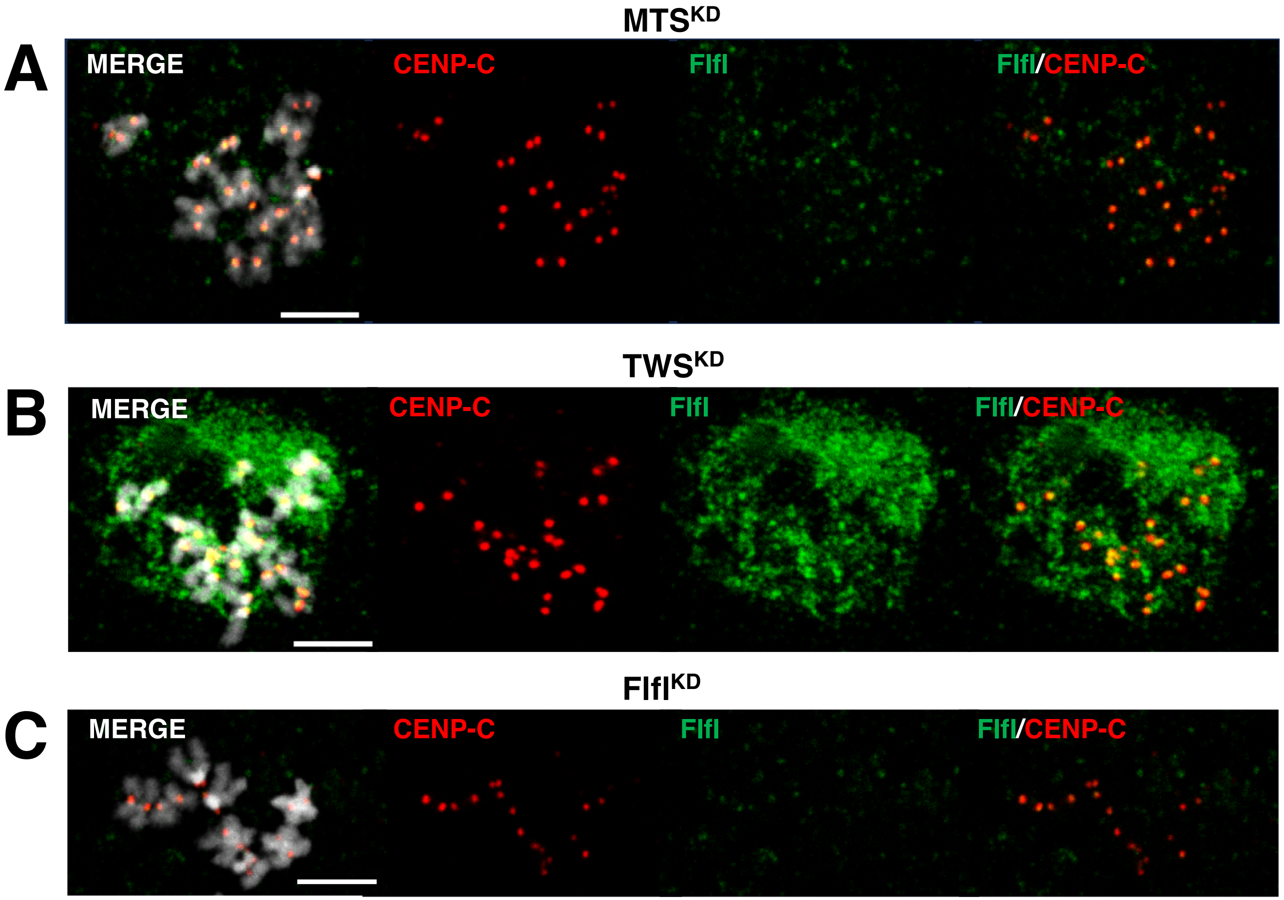
MTS^KD
A
MERGE
CENP-C
Flfl
Flfl/CENP-C
TWS^KD
B
MERGE
CENP-C
Flfl
Flfl/CENP-C
Flfl^KD
C
MERGE
CENP-C
Flfl
Flfl/CENP-C

**Figure S5**

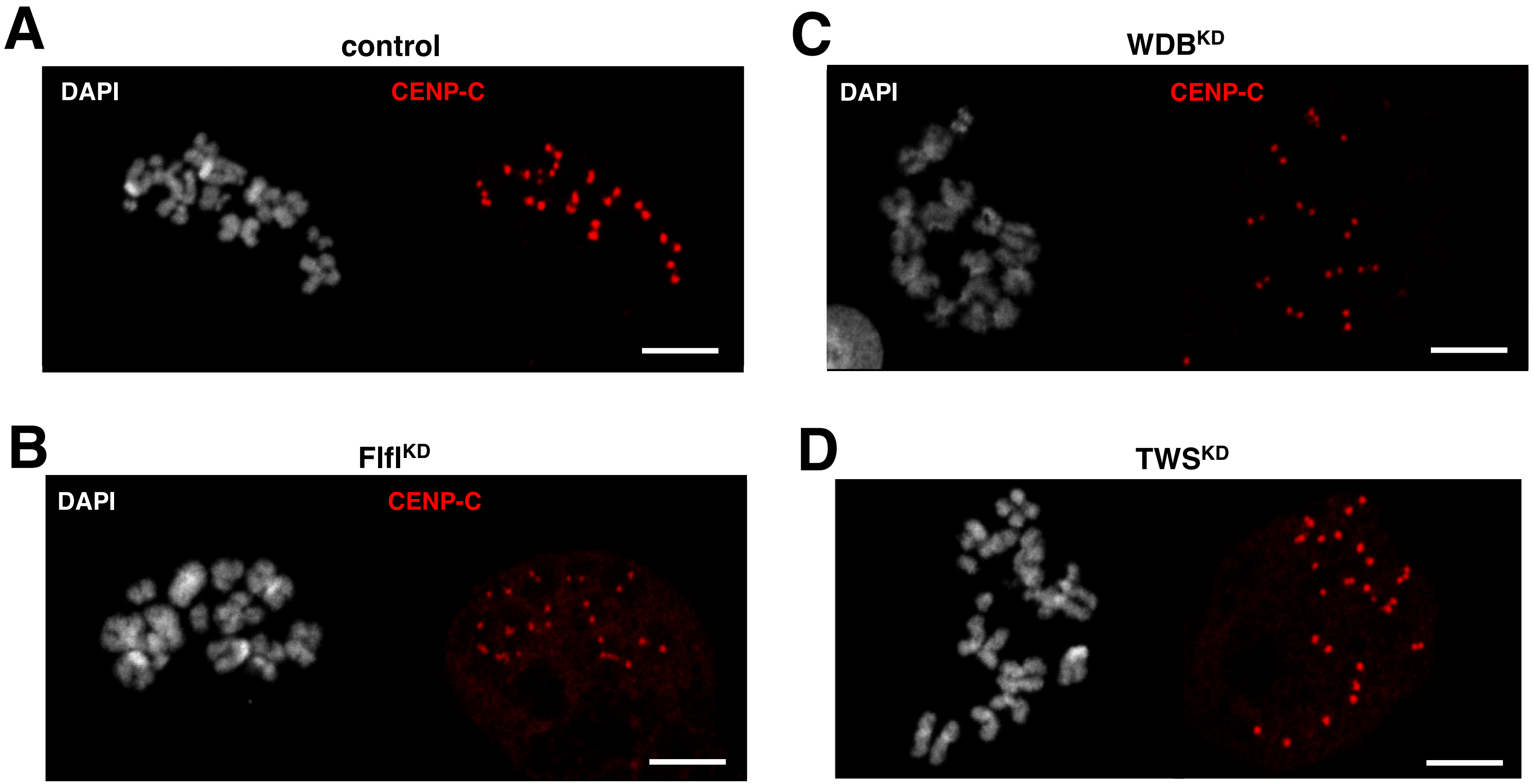
A
control
DAPI
CENP-C
B
Flfl$^{KD}$
DAPI
CENP-C
C
WDB$^{KD}$
DAPI
CENP-C
D
TWS$^{KD}$

Figure S6

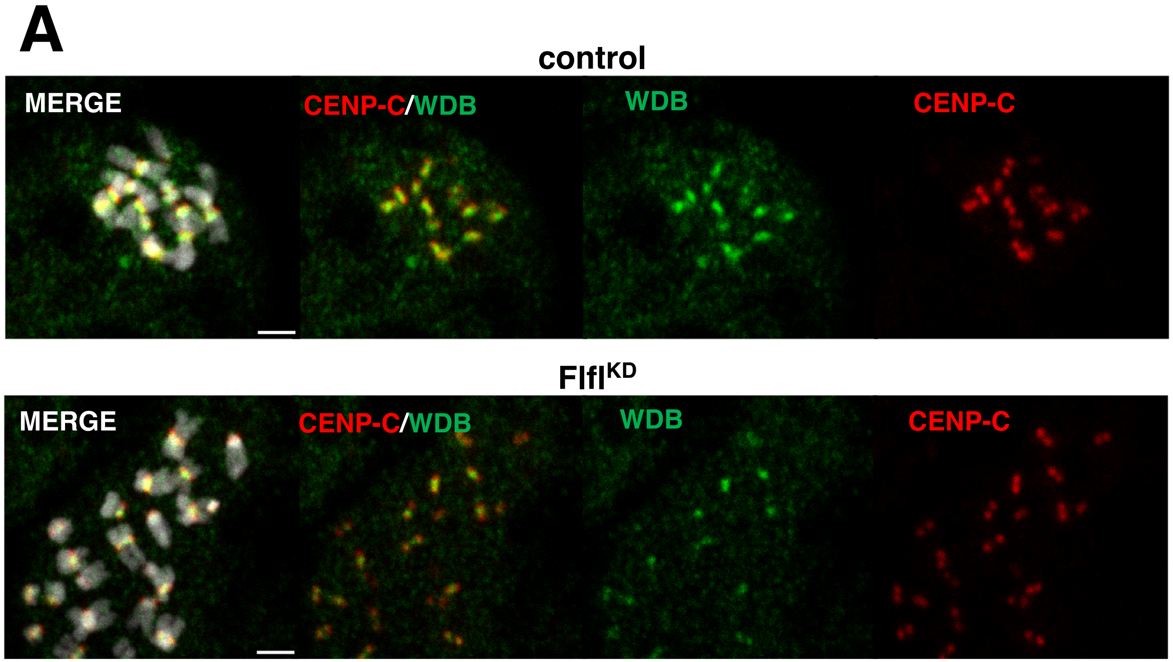
A
control
MERGE
CENP-C/WDB
WDB
CENP-C
FlflKD
MERGE
CENP-C/WDB
WDB
CENP-C

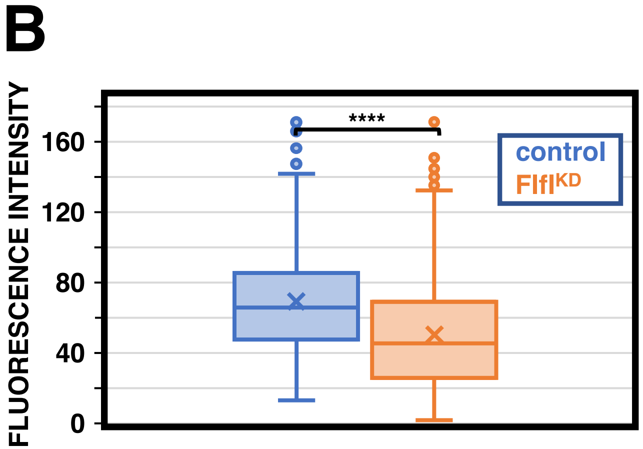
B
FLUORESCENCE INTENSITY
160
120
80
40
0
****
control
FlflKD

Figure S7
